# Evaluation of the Utility of a Research Ethics Training Course to Graduate Students

**DOI:** 10.1101/2025.09.22.677745

**Authors:** Michael D. Schaller, Peter H. Mathers

## Abstract

Concerns about the integrity of scientific research and the erosion of public trust in science led to policy recommendations to improve the responsible conduct of research. One recommendation was to increase scientific integrity through training, and numerous funding agencies mandated training in the responsible conduct of research for graduate students and postdoctoral fellows. Many institutions implemented training on consensus recommended topics. This study was initiated to evaluate graduate student perceptions of the utility of training in the responsible conduct of research during their dissertation research. The study was survey-based and captured the responses of first-year doctoral students enrolled in a semester-long course in the responsible conduct of research and past participants in the course. Students enrolled in the course demonstrated a gain in knowledge following training and an increase in self-efficacy in their ability to handle issues in scientific ethics. These students could envision how most of the topics in the course could be beneficial as they progressed in their scientific careers. Past participants in the course were asked to reflect upon their graduate studies and evaluate the impact of training in the responsible conduct of research. Many respondents identified sessions in the course that were useful at this early career stage and lessons learned from the course helped them navigate or avoid ethical challenges in their research. The results demonstrate that students appreciate the value of training in research integrity and that they are applying concepts from their training very early in their careers.

---

Research integrity and the responsible conduct of research (RCR) are essential for the continued advancement of science and the development of solutions to emerging problems impacting human health and innovative technologies to improve lives. The progress of research is incremental whereby current research builds upon the foundation of past research to advance the field. Integrity and RCR are necessary for members of the scientific community to trust the results from other researchers to build upon their findings rather than unnecessarily duplicating their studies and slowing advancement of the field (L. Bouter 2024). Sustained federal support for research since the second world war led to the rise of America’s scientific enterprise and is directly responsible for countless breakthroughs and technological advances. Continued support to drive scientific discovery depends upon the public trust that scientific research is performed with integrity and in the public interest.

Reports from several US national committees have stressed the importance of research integrity and the responsible conduct of research and provided broad recommendations to enhance the ethical performance of research (National Research Council 2002; National Academies of Sciences Engineering and Medicine 2017). U.S. federal agencies were encouraged to support studies into scientific integrity to define underlying factors influencing the integrity of scientists and best practices to promote ethical conduct. Academic and research institutions were encouraged to establish a culture of scientific integrity by implementation of comprehensive policies designed to promote and sustain the ethical performance of research. One strategy to comprehensively address the issue is the adoption of a Research Integrity Promotion Plan that is linked to codes of conduct for research. This should include the institutional goals and guidelines, and procedures and support systems to facilitate accomplishing their goals (L. Bouter 2020). One facet of the recommended policy approach was the development of education and training for all scientists that will promote the integrity of research (National Research Council 2002; National Academies of Sciences Engineering and Medicine 2017). Traditionally, training in all aspects of research was the responsibility of the scientific mentors of trainees. Research at the time of these early reports suggested that insufficient education in research integrity was provided for the trainees by their mentors (Eisen and Berry 2002; M. Kalichman 2013a). Further, scientists believe training in integrity in some areas of research should be the responsibilities of mentors, while the responsibility for training in other areas lies with the institution (Eisen and Berry 2002; Titus and Ballou 2014). Thus, the recommendations were to build scientific integrity training into the curriculum and embed training longitudinally and across different components of graduate student and postdoctoral training.

U.S. federal funding agencies mandated training in the responsible conduct of research for all trainees supported by agency funds (Steneck and Bulger 2007; M. Kalichman 2013a). The NIH provided the first guidelines of topics to be included in training and these guidelines have been modified over time (National Academy of Engineering 2013). However, federal agencies provided little additional guidance for implementation of training and assessment of the effectiveness of training. Academic institutions implemented training programs to meet the initial mandates and have expanded training broadly to trainees regardless of their source of funding (Resnik and Dinse 2012). However, given limitations on guidance for implementation, there has been considerable variability in RCR training programs across institutions. Training programs can vary across all aspects including goals, delivery, duration, content, methods and evaluation (M.W. Kalichman 2007; M.W. Kalichman and Plemmons 2007; Mulhearn et al. 2017; Watts et al. 2017). Further, the outcomes expected and measured differ between programs. Clearly, the intent of training in scientific integrity is to reduce occurrences of scientific misconduct, which are egregious, but also to reduce occurrences of questionable research conduct, which are less egregious but undermine the reproducibility of research and are likely driven by the incentives in academic research (Steneck 2006; L. Bouter 2020). Behavioral outcomes (like ethical actions) are difficult to measure and difficult to tie back to specific interventions. Other outcomes that can be evaluated more directly and are indicators of future ethical behavior are more realistic measures. These include the measurement of attainment of knowledge regarding guidelines for research and misconduct, skills to approach ethical problems, attitudes toward ethical issues, sensitivity/awareness of arising ethical issues and judgement (Resnik 2014; McGee 2014; M.W. Kalichman and Plemmons 2015). Early analyses of the outcomes of training in RCR revealed little or modest effects (Antes et al. 2009; Antes et al. 2010), while more recent analyses suggest training is having a larger impact on trainees (Watts et al. 2017; Abdi et al. 2021). Deeper analyses of these studies have identified types of training, instructional methods, and type of curriculum that are most effective in training in RCR (Todd et al. 2017; Torrence et al. 2017; Mulhearn et al. 2017). These are useful resources for the design or modification of training programs in scientific integrity.

While many studies have been designed to measure the effectiveness of RCR training, particularly training of graduate students, few have addressed the usefulness of training to those students. Such insight will be valuable for modifications to improve curriculum content and reception of training by students. In this study, the utility of RCR training was assessed from the graduate student perspective. Student attitudes, self-efficacy and knowledge were measured in two cohorts of trainees enrolled in an RCR training course for first-year graduate students. Trainees were asked their perception of the usefulness of different RCR sessions for their future research careers. To complement these findings, past trainees who participated in this training program were asked in hindsight how useful different sessions were for their dissertation research. The findings suggest that graduate students appreciate the purpose of RCR training and utilize lessons learned from these sessions in their graduate career.

## Methods

### Courses included

A year-long series of two courses in scientific ethics has been taught for over 15 years at our institution. The courses included are a fall semester course entitled, BMS700 Scientific Integrity, and a spring semester course entitled, BMS701 Scientific Rigor and Ethics. Each course is comprised of seven 2-hour sessions spread across each semester covering consensus topics (**see Online Resource 1**)(Steneck 2007; Institute of Medicine 2009; M. Kalichman 2013b). The course has evolved, for example by incorporation of the film “Picture a Scientist” (Cheney and Shattuck 2020) and “The Lab: Avoiding Research Misconduct” exercise from the Office of Research Integrity at the NIH (Office of Research Integrity US Department of Health and Human Services) in replacement of sessions on entrepreneurship and mentoring. At our institution, several other important RCR topics, including mentor/mentee relations, are covered in a professional skills course. Each session of BMS700/BMS701 begins with small group discussion of a short pre-read article. A faculty/staff member makes a didactic presentation and facilitates discussion of the topic. The course co-coordinators (the authors) also participate in facilitating discussion. Each session ends in small group discussion about several case studies, followed by a general group discussion. Students participating in these scientific ethics courses primarily included those enrolled in the Biomedical Sciences Graduate Programs, along with others from various departments across our campus. Specific demographics of the programs included can be found in the Results section.

### Demographics collection

De-identified demographics of all students enrolled in BMS700 or BMS701 from fall 2017 through fall 2024 were obtained from the Registrar’s Office. A survey was designed to capture the demographics of the respondents to both in-class and online surveys. Some of the questions on this survey captured different data than available from the Registrar’s Office.

### Survey types

Two independent surveys were constructed and approved by our Institutional Review Board (Protocol #2308831247 and #2306805348). The first survey was an in-class survey performed in a pre-test/post-test format during the fall semesters of 2023 and 2024. Here, the students were asked for demographic data (part 1) (only in fall 2024) and a combination of an attitude assessment of ethics training and their self-efficacy in managing scientific ethical issues, along with skills-based questions to gauge short-term effectiveness of the ethics training across the semesters (part 2) (**see Online Resource 2**). Students were asked to create a self-selected personal identification number to use on all surveys in the course, allowing answers to different surveys to be linked in an anonymous fashion. The second of these surveys was an email-solicited, online survey that was distributed to previous classes. This study was a retrospective analysis of students who already had completed the course. They were asked for demographic data (part 1) and coursework assessment data to gauge long-term utility of the ethics training (part 2) (**see Online Resource 2**). Due to the evolution of the course, a minor part of the content differed for these students and was reflected in the survey questions. The survey was open for a month. Students received three reminders to complete the survey and were incentivized with a random drawing for two $25 gift cards. Students wishing to participate in the draw submitted a self-selected PIN and the draw was managed by a member of staff in the Department of Biochemistry and Molecular Medicine. In both surveys, the demographics and assessment surveys were unlinked to preserve anonymity. This was critical since our sample size and institutional demographics would allow identification of some students based upon demographic responses.

### Survey participants

The surveys of current students were administered in class after requesting informed consent. Approximately 94% of students consented to participate in the study. Survey data was included for all students whose self-selected PIN could be matched on the pre- and post-surveys. For the fall 2023 class, the survey responses of 38 students were included and for the fall 2024 class, responses of 45 students were included.

Potential participants in the online survey included graduate students (mostly doctoral but some Master’s students, as well) in the biomedical and biological sciences, along with others who required scientific ethics for F- and K-awards. This group encompassed classes ranging from the fall semester of 2017 to spring semester of 2024, including the intervening classes. Emails requesting participation in the retrospective surveys were sent to all formerly enrolled students (n = 326). From this list, 41 email addresses were no longer active. Therefore, 285 students were invited to participate in the retrospective survey. The response rate was 26.0% (n = 74) for the assessment survey of former students. All participants consented to inclusion of their anonymous data for this analysis.

### Statistical Analyses

The distribution of the scores on the pre- and post-tests for knowledge was Gaussian. Statistical analysis was performed using a two-tailed, paired t-test. Ordinal values were used for the Likert responses. When the five questions on self-efficacy were summed, the distribution of the data was Gaussian, therefore a two-tailed, paired t-test was performed. Results were judged to be statistically significant if p < 0.05. Effect sizes were determined by calculating Cohen’s d. The distribution of scores in many of the individual self-efficacy questions were not Gaussian and were ordinal. Therefore, these data were analyzed using a two-tailed Wilcoxon pairs signed rank test using Pratt’s Method to handle ties. To assess consistency of responses on the online survey, responses were divided between two cohorts based upon the performance of a particular type of research or identification of a specific session as useful in helping navigate or avoid an ethical situation. For example, students performing research on human subjects formed one cohort, and students not performing research on human subjects formed a second cohort for analysis of the question addressing usefulness of training in human-subjects research. The distribution of many of these responses was non-Gaussian, and the scale was ordinal. A two-tailed Mann Whitney U test was performed for comparison of these cohorts. Statistical analyses were performed using GraphPad Prism or Python.

### Internal Survey Validation

To assess construct validity, an inter-item correlation analysis was performed on the post-test survey questions about self-efficacy and on the survey questions related to attitudes. As the results are non-Gaussian, Spearman r was calculated for each pair of questions. The results of the self-efficacy questions show a Spearman r value greater than 0.358, with the exception of the correlation between the question about knowledge and whether the respondent could apply ethics, which was 0.262 (**Table 1**). The internal consistency reliability for the self-efficacy questions was determined by calculating Cronbach’s alpha (0.8134). These results suggest convergence of these survey questions, i.e. that they are measuring related concepts. Similar results were found for the attitudes survey questions. Spearman r values were greater than 0.5 for each pair of questions, except for the question about frequency of misconduct, which was less than 0.25. Cronbach’s alpha was 0.8619. If the frequency of misconduct question was excluded, Cronbach’s alpha was 0.916. These results suggest convergence of the results of the attitudes questions. To measure divergence, an intra-construct correlation analysis was performed comparing the sum of the responses to the self-efficacy questions and the sum of the responses of the attitudes question (excluding the question about frequency of misconduct). Spearman r was 0.182 reflecting a low correlation, suggesting that the two constructs were measuring different concepts. These results support the validity of the in-class survey.

**Table 1.** Internal Validation of Surveys.

| **Inter-Item Correlation of Responses to Self-Efficacy Questions (Spearman $r$)** |  |  |  |  |  |
| --- | --- | --- | --- | --- | --- |
|  | Knowledge | Handle Issues | Find Resources | Maintain Ethical Beh | I can apply ethics |
| Knowledge | 1 |  |  |  |  |
| Handle Issues | 0.583 | 1 |  |  |  |
| Find Resources | 0.522 | 0.605 | 1 |  |  |
| Maintain Ethical Beh | 0.481 | 0.479 | 0.526 | 1 |  |
| I can apply ethics | 0.262 | 0.404 | 0.396 | 0.358 | 1 |
| **Inter-Item Correlation of Responses to Attitudes Questions (Spearman $r$)** |  |  |  |  |  |
|  | Training is important | Training prevents misconduct | Ethics is important | Frequency of misconduct |  |
| Training is important | 1 |  |  |  |  |
| Training prevents misconduct | 0.654 | 1 |  |  |  |
| Ethics is important | 0.652 | 0.509 | 1 |  |  |
| Frequency of misconduct | 0.232 | 0.191 | 0.241 | 1 |  |

## Results

### Demographics

Initial training in the responsible conduct of research for biomedical science graduate students at our institution occurs in two in-person courses in the fall and spring semester of their first year (**see Online Resource 1**). Graduate students in other programs who wish to take or require a scientific integrity course often take one or both of these courses. The demographics of students registered for these courses from the fall semester 2017 through fall semester 2024 are summarized in **Table 2**. In-class surveys were performed in the fall semester course in 2023 and 2024 since the fall semester course has a larger enrollment from a broader range of programs. In the fall 2023 and 2024 semesters, there were 101 registered students combined, with 59.4% from the biomedical sciences, whereas the spring semester 2024 and 2025 courses had 64 registered students (mostly a subset of participants in the fall course) with 87.5% from the biomedical sciences. A demographic survey was administered to the class in fall 2024 (**Table 2**).

**Table 2.** Student Demographics.

|  | Students 2017-2024 | Fall 2024 | Respondents to Online Survey |
| --- | --- | --- | --- |
| **n** | 334 | 48 | 85 |
| **Gender** |  |  |  |
| Male | 37% | 20.8% | 27.1% |
| Female | 63% | 72.9% | 65.9% |
| Non-binary | n/a<sup>a</sup> | 2.1% | 3.5% |
| Transgender | n/a | 0% | 1.2% |
| Prefer not to answer | n/a | 4.2% | 2.4% |
| **Ethnicity** |  |  |  |
| American Indian/Alaskan Native | 0.6% | 0% | 0% |
| Asian | 19.2% | 18.8% | 20% |
| Black/African American | 5.4% | 8.3% | 3.5% |
| Hawaiian/Pacific Islander | 1.2% | 0% | 0% |
| Hispanic | 2.4% | 2.1% | 0% |
| White | 74.9% | 58.3% | 71.8% |
| Middle Eastern | n/a | 16.7% | 3.5% |
| Self-describe | n/a | 0% | 1.2% |
| **Citizenship** |  |  |  |
| US Citizen | 73.4% | 64.6% | 76.5% |
| Permanent Resident | 4.2% | n/a | n/a |
| Non-Immigrant | 22.5% | 35.5% | 22.4% |
| **First Generation College** |  |  |  |
| Yes | n/a | 52.1% | 63.5% |
| No | n/a | 47.9% | 36.5% |
| **Self-Identify Disability** |  |  |  |
| Yes | n/a | 25% | 17.6% |
| No | n/a | 68.8% | 82.4% |
| Prefer not to answer | n/a | 6.25% | 0% |
| **Socioeconomic Disadvan.** |  |  |  |
| Yes | n/a | 37.5% | 41.2% |
| No | n/a | 45.8% | 57.6% |
| Prefer not to answer | n/a | 16.7% | 1.2% |
| **Program** |  |  |  |
| Biomedical Sciences | 77.0% | 63.0%<sup>b</sup> | n/a |
| Biology | 12.3% | 24.1%<sup>b</sup> | n/a |
| Public Health | 6.3% | 3.7%<sup>b</sup> | n/a |
| Pharmacy | 2.7% | 0%<sup>b</sup> | n/a |
| Engineering | 1.2% | 9.3%<sup>b</sup> | n/a |
| Other | 0.6% | 0%* | n/a |
| a - n/a – question/response was not included; b – n =54 for program data |  |  |  |

### Motivation to take the course

Students were asked why they took the course and 56.6% reported taking it because it was mandated, 33.7% responded that it was their responsibility and 9.6% took the course out of interest. To determine if their motivation to take the course impacted performance, attitudes or self-efficacy, results were compared between the cohort of students taking the course because it was mandated and the cohort taking the course for another reason. Statistical analysis revealed no significant difference in the responses to questions related to attitude, self-efficacy or knowledge between the two cohorts (**Online Resource 3**). The results demonstrate that motivation to take the course did not impact the outcomes measured.

### Students Learned

Two measurements were made with the in-class surveys to determine if the students learned. First, a survey following each session asked if they learned important information on this topic and the answers were recorded on a Likert scale (1 to 5). A large majority of students reported learning in each session (see **Fig. 1a**). The median Likert score for three of the sessions was 4 and the median score for the other four sessions was 5. Second, the students took a pre-test in the first class and a post-test in the final class to measure changes in knowledge. Out of 14 questions, the average number of correct questions on the pre-test was 6.9 +/- 2.1 and the average on the post-test was 9.5 +/- 2.08 (see **Fig. 1b**). Individual pre-tests and post-tests could be linked due to an anonymous, self-selected unique identifier chosen by each student allowing comparison of individual performances between the two tests. The majority of students performed better on the post-test (80.7%). The distribution of the scores was Gaussian, therefore the differences were analyzed using a two-tailed, paired t test (n = 83, t = 10.51, df =82, p < 0.0001, d = 1.257). Both measures demonstrate that the students gained knowledge from the course.

**Fig. 1.**
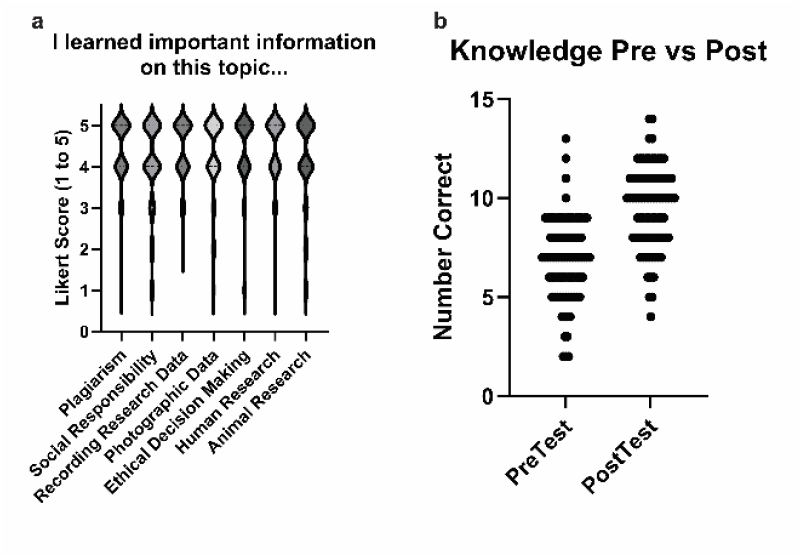
The students learned. **a)** In a survey following each of the 7 sessions, students were asked to select a response on the Likert scale (1 = strongly disagree; 5 = strongly agree) to the statement “I learned important information about this topic”. Their responses are shown in the violin plot. The median for 4 sessions was a score of 5 and the median for 3 sessions was a score of 4. **b)** Student knowledge was independently assessed on a pre-test and post-test consisting of the same 14 questions (2 from each session). The number of correct answers on the pre-test and post-test for each student is shown. Pre-test mean +/- sd = 6.9 +/- 2.1, 95% CI = 6.45-7.36. Post-test mean +/- sd = 9.5 +/- 2.08, 95% CI = 9.08-9.98. Student performance on the pre- and post-test was analyzed using a two-tailed, paired t test (p < 0.0001, d = 1.257).

### Student Attitudes

Students were asked questions to assess their attitudes about training in scientific ethics and how frequently they thought misconduct occurred (**Fig. 2**). Median Likert scores to survey prompts asking the importance of training, whether training prevents misconduct and if ethics is important for scientific inquiry on the pre-tests and post-tests were 5 (i.e. strongly agree). The responses to these questions were convergent (Cronbach’s alpha = 0.916). Improvement in attitudes was anticipated following the course, but the positive attitudes demonstrated on the pre-test precluded measuring a change in attitude. It was anticipated that stratification based upon attitudes presented on the pre-test might provide insight into the role of attitude in engagement in each session, learning and increasing self-efficacy by the end of the course. The uniformly high scores on questions related to attitude on the pre-test did not allow stratification to perform this analysis. With regard to student perceptions of the frequency of science misconduct, a large majority believe misconduct occurs occasionally or frequently (**Fig. 2e**). A similar distribution of responses was found on the pre-test and post-test.

**Fig. 2.**
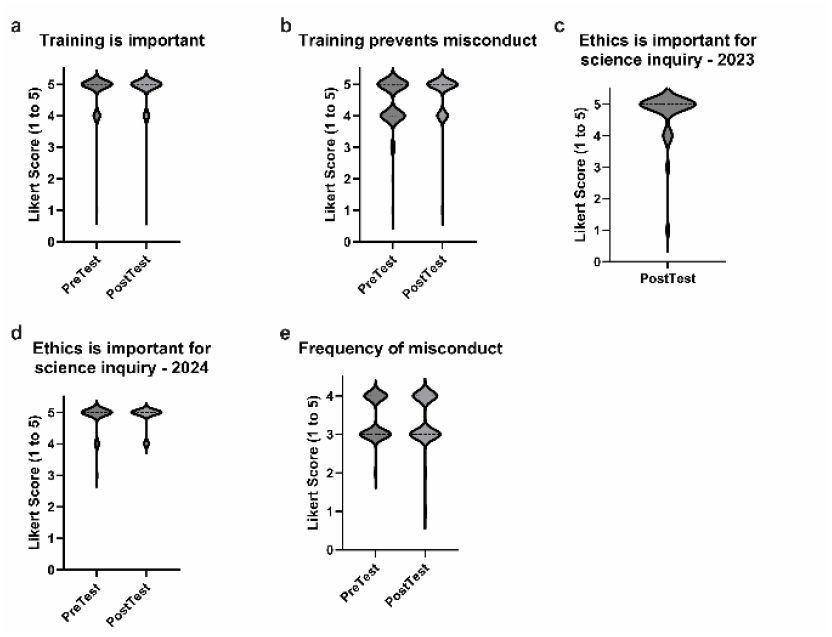
Student attitudes. Student attitudes about research ethics were assessed by their response on a Likert scale (1 = strongly disagree to 5 = strongly agree) to three statements. Attitudes were captured on a pre-test and post-test. **a)** Training is important (fall 2023 and 2024 combined), median = 5 on both tests. **b)** Training prevents misconduct (fall 2023 and combined), median = 5 on both tests. **c)** Ethics is important for scientific inquiry, post-test results for Fall 2023 (the question was inadvertently omitted from the pre-test), median = 5. **d)** Ethics is important for scientific inquiry, pre- and post-test results for Fall 2024, median = 5 for both tests. **e)** Student perception of the frequency of scientific misconduct was measured on a 4-point scale (1 = never, 2 = rarely, 3 = occasionally, 4 = often), median = 3.

### Increased Self-Efficacy

Changes in student self-efficacy were measured in two ways. First, on the post-test, students were directly asked if the course increased their ability to handle issues in scientific ethics (**Fig. 3a**). The median score on a Likert scale (1 = strongly disagree; 5 = strongly agree) was 4. Therefore, the students perceived an increase in self-efficacy over the duration of the course. Second, the students were asked to respond to five survey questions related to self-efficacy on a pre-test and post-test. Differences in scores between the two tests were used as a measure of the change in self-efficacy from the beginning to the end of the course. The sum of the scores of the five questions was used as one measure of changes in self-efficacy (**Fig. 3b**). The average summed score on the post-test was 2.24 points higher (reflecting a 14.5% increase in score over the pre-test). Individual student scores on the pre-test and post-test were analyzed using a two-tailed, paired t test (p < 0.0001, d = 0.9233). This result also supports an increase in self-efficacy during the course. Pre- and post-test responses to each individual self-efficacy question were also analyzed (**Fig. 3c-g, Table 3**). The responses to these questions were convergent (Cronbach’s alpha = 0.8619 – performed on the post-test). Scores increased on the post-test for responses to prompts about the student’s confidence in knowledge, ability to handle ethical issues, in finding resources to help decision making, and maintaining ethical behavior (**Fig. 3c-f**). The results of two-tailed Wilcoxon matched pairs signed rank tests (using Pratt’s method to handle ties) demonstrated a significant difference between the pre- and post-test for these four questions (**Table 3**). The responses to the firth question, student confidence that they can apply ethics to make decisions, were high on both the pre-test and post-test (median = 4) (**Fig. 3g**). There was no statistical difference in responses between the pre- and post-test for this question (**Table 3**). In toto, these data demonstrate that self-efficacy increased from participation in the course.

**Table 3.** Statistics for Changes in Self-Efficacy (by the question)

| Question | # pairs | # ties | Sum of signed ranks (W) | p value |
| --- | --- | --- | --- | --- |
| Knowledge about ethical issues | 83 | 28 | 2542 | <0.0001* |
| Ability to handle ethical issues | 83 | 36 | 1916 | <0.0001* |
| Confidence to find resources | 83 | 24 | 2652 | <0.0001* |
| Confidence in maintaining ethical beh. | 82<sup>a</sup> | 43 | 846 | 0.035* |
| I can apply ethics to make decisions | 83 | 50 | 603 | 0.1628 |
| a - One student did not answer this survey question on the pre-test; * $p < 0.05$ | | | | |

**Fig. 3.**
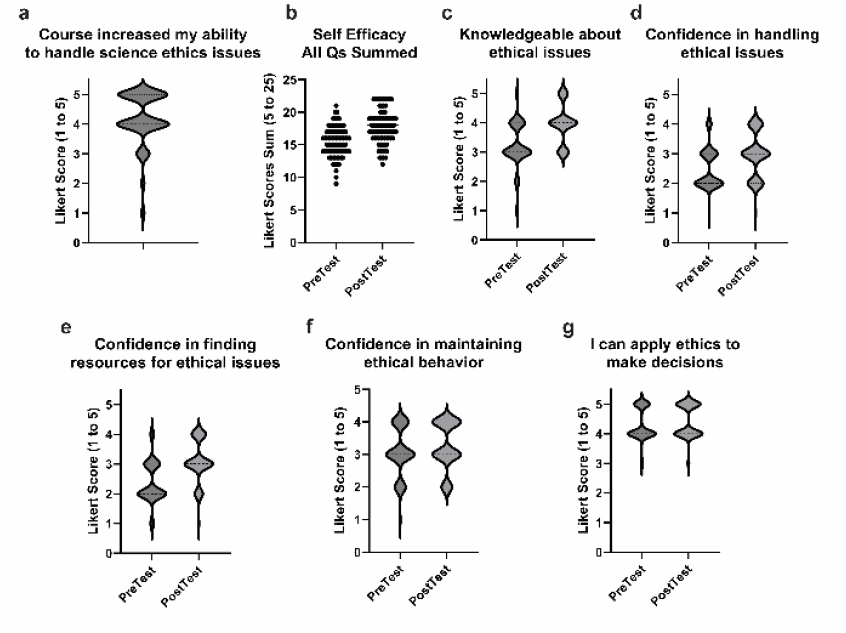
Students’ self-efficacy. Student self-efficacy was measured by survey responses on a Likert scale (1 = strongly disagree, 5 = strongly agree) and are illustrated in violin plots. **a)** A post-test survey question asked the students if the course increased their ability to handle ethical issues, median = 4. **b)** The responses to five survey questions related to self-efficacy were summed. Comparison of the sums on the pre-test with the sums on the post-test indicate that students’ self-efficacy increased, pre-test mean +/- sd = 15.46 +/- 2.3, 95% CI = 14.96 to 15.96; post-test mean +/- sd = 17.7 +/- 2.55, 95% CI = 17.14 to 18.26. The data is Gaussian and was analyzed with a two-tailed, paired t test (p < 0.0001, d = 0.9233). **c-g)** Pre- and post-test responses to five individual questions related to self-efficacy are plotted. **c)** Pre-test median = 3; post-test = 4. **d)** Pre-test median = 2; post-test = 3. **e)** Pre-test median = 2, post-test = 3. **f)** Pre-test median = 3; post-test = 3. **g)** Pre-test median = 4; post-test = 4.

### Student Perceptions of Utility of Training in Different Topics

Student perceptions of the utility of training in each topic were measured with a survey question in each session asking if the students could see the future utility of training. The results of the survey indicate that the students overwhelmingly see the utility of the training that was provided, with a median score of 5 on a 5-point Likert for six of the sessions and a median score of 4 for the session on animal research (**Fig. 4**).

**Fig. 4.**
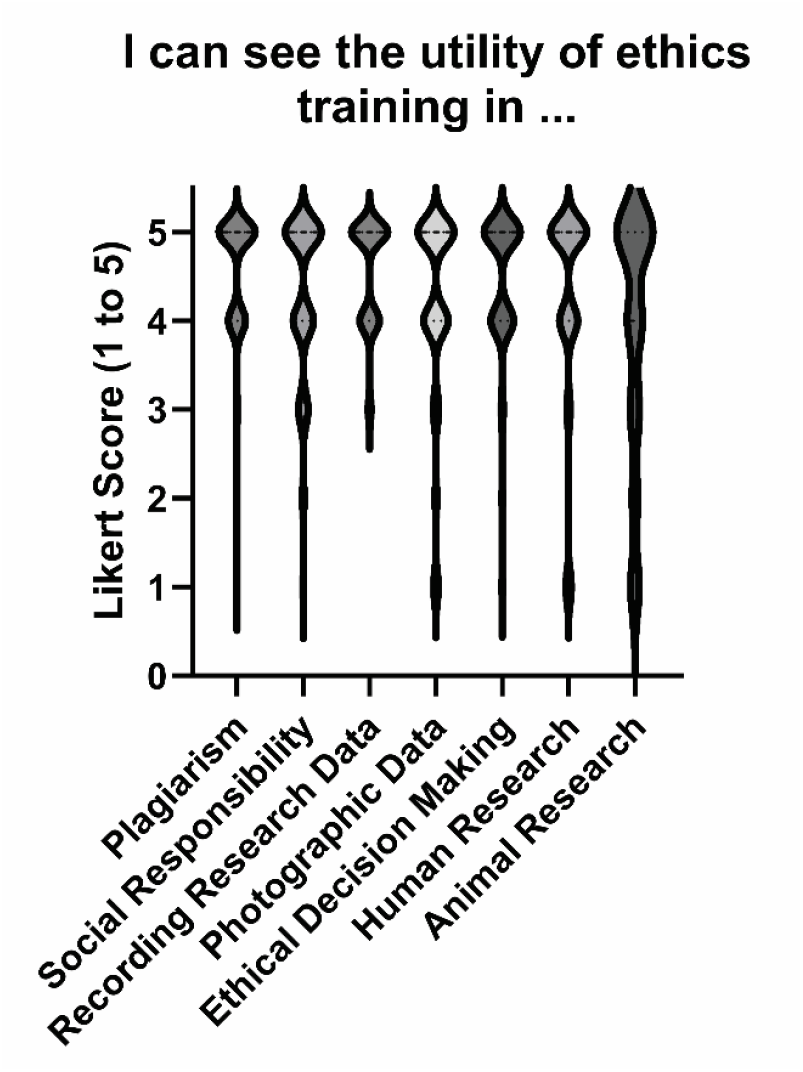
Students views on the utility of ethics training. Results of a survey response from each session measuring if the students could see the utility of the topic at the time of training. The median score for all sessions was 5, except for the session on animal research, which had a median score of 4.

### Student Experiences Following Initial Training in Scientific Ethics

In order to assess the perceived utility of RCR training in the dissertation work of graduate students, an online survey was distributed to all students who were enrolled in the BMS700/BMS701 courses since Fall 2017. Emails soliciting participation in this study were sent to 285 past participants in the course with a known active email address. Responses were received from 74 individuals (26%). The demographics of respondents is included in Table 2. The distribution of respondents along their career path is illustrated in **Fig. 5a**. The most responses were received from students who had already completed their graduate studies or were in Year 3 or 4 of graduate school. Of these respondents, 44.6% reported additional RCR training. These included other courses, online training, departmental and T32 training grant offerings, seminars and Society/Conference workshops (**Fig. 5b**). Despite these additional training opportunities, the survey focused the respondents on the content and experiences from BMS700/BMS701 and asked how these helped navigate different situations.

**Fig. 5.**
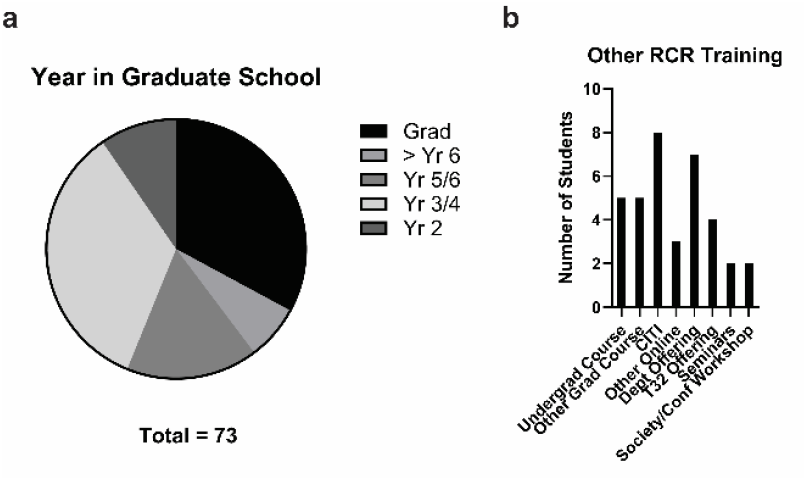
Online Survey Respondents and Other RCR Training. **a)** Survey respondents were asked to identify their current year in their graduate program. **b)** Approximately 45% of respondents reported additional training in RCR. The types of additional training reported and the number of students reporting each type of additional training are illustrated.

The survey asked respondents about the utility of training in the BMS700/BMS701 courses with questions related to specific topics. For example, the survey asked for a response on the Likert scale (strongly disagree = 1 to strongly agree = 5) to the statement “I have used lessons from one or more sessions of the course for managing my lab notebook or data collection”. Four topics were relevant to all of the respondents: data management, mentor relations, plagiarism and management of issues related to scientific misconduct. Other topics were specifically relevant to a subset of respondents, e.g. the usefulness of training in human subject research is most relevant to students performing research involving human subjects. For these topics, the responses of the subset of respondents to whom the topic was most relevant is presented.

The responses to questions relevant to all respondents are illustrated in **Fig. 6a**. The median Likert score for each of these questions was 4. Seventy percent of the respondents published at this point in their graduate career and another 10.8% had submitted but not yet published a manuscript. The responses of these students to the question about usefulness of the authorship session are shown in **Fig. 6b** (median Likert score = 4). Fifty-eight percent of respondents have participated in peer review. Their responses to the utility of the peer review session are shown in **Fig. 6c** (median Likert score = 4). The responses of the seventeen respondents who conducted research using human subjects to the usefulness of the sessions on human subject research are shown in **Fig. 6d** (median Likert score = 4). The research of two-thirds of the students used animals, and **Fig. 6e** shows their responses about the usefulness of RCR training on animal research in the course (median Likert score = 4). Thirty-seven percent of the respondents report performing collaborative research during their dissertation research. Their responses related to the utility of the session on collaborative research are illustrated in **Fig. 6f** (median Likert score = 4). Only 19% of the students reported managing a conflict of interest. Their responses to the question of usefulness of conflict-of-interest training are shown in **Fig. 6g** (median Likert score = 4). These results demonstrate that participants in the major graduate-level RCR course at our institution retrospectively found the training useful during their tenure as a graduate student.

**Fig. 6.**
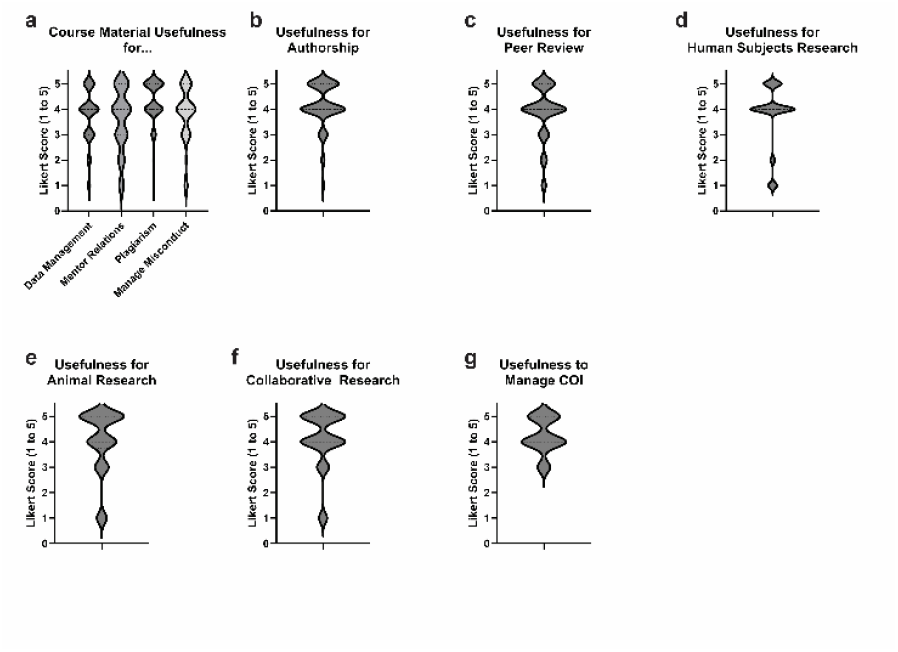
Online Survey Respondents Report the Usefulness of RCR Training in Graduate School. Survey responses to a query about the students’ retrospective view of the usefulness of individual RCR training sessions in BMS700 and BMS701 are illustrated. **a)** The survey results of all respondents to the questions about data management, mentor relations, plagiarism and managing misconduct are reported (n = 74). Note that the plagiarism question was flipped and was phrased “The plagiarism session did NOT provide useful examples to avoid ethical issues”. The responses were inverted for depiction in panel A. **b)** Survey responses from respondents who have published or submitted a manuscript regarding the usefulness of the session on authorship are shown (n = 60). **c)** Survey responses from respondents who have performed peer review regarding the usefulness of the session on peer review are illustrated (n = 43). **d)** Reponses of the 17 students who performed human subjects research regarding the usefulness of the training session on human subjects are shown. **e)** The responses of 50 students who performed animal research regarding the usefulness of the training session on animal research are illustrated. **f)** The usefulness of training in collaborative research as reported by respondents who have participated in collaborative research are shown (n = 27). **g)** Fourteen students report managing a conflict of interest. The responses about the usefulness of the training session on conflict of interest is illustrated. The median response for all 10 of these survey questions was 4.

Two additional survey questions were asked to corroborate the responses to queries probing the utility of different sessions. The first question asked if there were topics in the BMS700 or BMS701 courses that were useful in helping to navigate specific situations. Twenty-eight respondents (37.8%) said that none of the sessions helped navigate situations. The other 62% of respondents provided 151 responses identifying a session that was useful (**Fig. 7a & b**). The second question asked if there were topics in the courses that were useful to help avoid unethical situations. Only 17 respondents (23.6%) said that none of the sessions helped them avoid unethical situations. The other 76% of respondents provided 250 responses identifying a session that was useful in avoiding unethical situations (**Fig. 7c & d**). These results dramatically illustrate that graduate students draw upon their experiences in an in-person RCR training course to navigate ethical situations and perhaps more importantly to avoid unethical situations during the course of their graduate career.

**Fig. 7.**
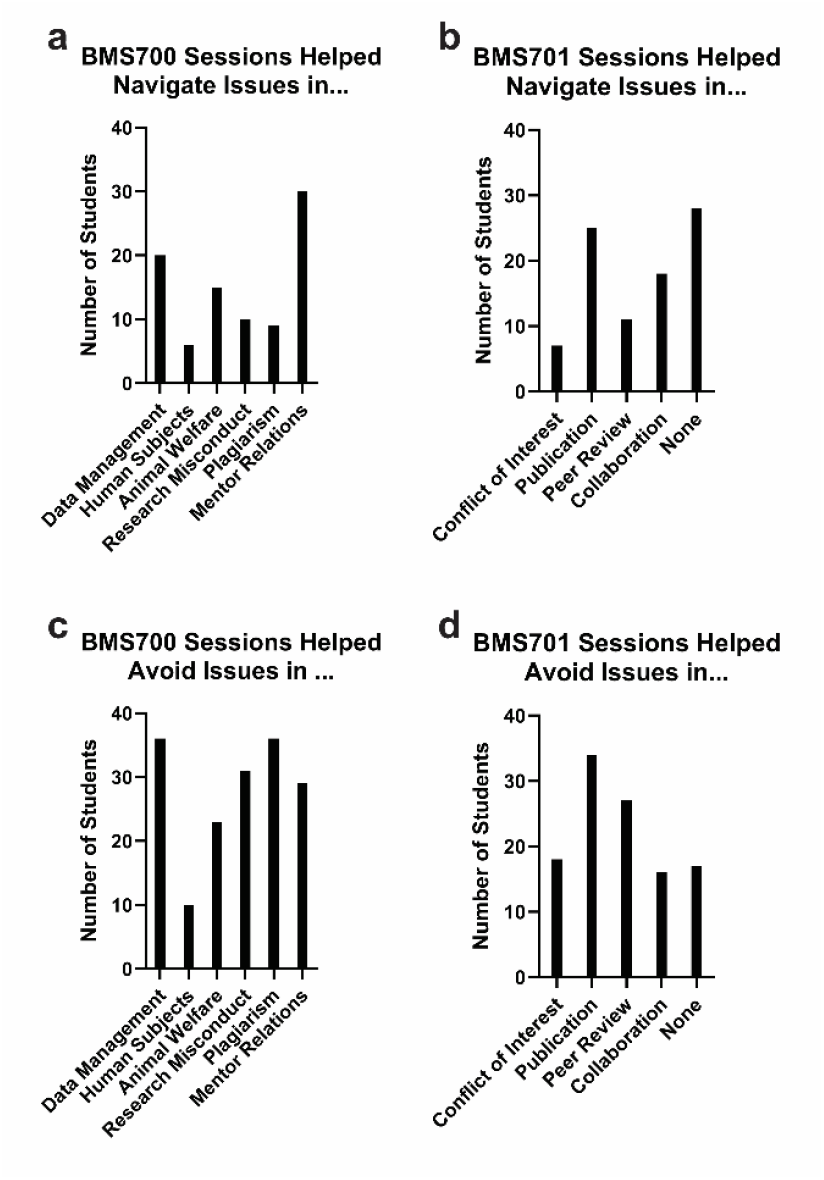
Student Identification of Sessions Used to Navigate or Avoid Ethical Issues. Students were asked to select training sessions that helped them navigate specific issues (**panel a & b**) or avoid ethical issues (**panel c & d**). Training sessions from the fall semester (BMS700) are shown in panels **a & c** and training sessions from the spring semester (BMS701) in panels **b & d**. The number of students reporting that none of the sessions helped them navigate or avoid ethical issues are included in panels **b & d**.

The survey data was further analyzed to establish internal consistencies in responses. There are two predictions for consistency, based upon the survey design. The first prediction is that individual sessions will be more useful to students performing research related to the session (e.g. human subjects research) or having an experience related to an individual session (e.g. authorship) than to students who do not perform a particular type of research or have an experience related to the session. The responses to the survey questions related to authorship, peer review, human subjects research, animal research and collaborative research were separated into two groups. Respondents who performed that type of research/had a relevant experience were assigned to one group, called the “yes” group, and the other group included the respondents who did not perform that type of research or did not have the relevant experience, called the “no” group. The Likert responses of the two groups were compared using a Mann Whitney U test to determine if the responses were statistically different. The responses between the two groups were statistically different for 4 out of 5 questions, with the exception being the question about the utility of RCR training in animal research (**Table 4**). The second prediction is that there will be concordance between the responses of the usefulness of a particular session, i.e. response to the question “I have used lessons from one or more sessions of the course for managing my lab notebook or data collection” (**Fig. 6**), and the sessions that the students identified that helped them navigate a situation or avoid an ethical issue (**Fig. 7**). Respondents were divided into two groups based upon their responses to the two questions about navigating a situation or avoiding an ethical issue, e.g. respondents selecting data management as a session that helped them navigate a situation or avoid an ethical issue were assigned to one group, called the “yes” group, and respondents who did not select data management as a session helping navigate or avoid an ethical issue were assigned to the other group, called the “no” group. The Likert scale responses of the two groups to the survey query “I have used lessons from one or more sessions of the course for managing my lab notebook or data collection” were compared using a Mann Whitney U test to determine if they two are statistically different (**Table 5**). The responses to 5 of the 9 questions were significantly different between the two groups. The results of these two analyses demonstrate consistency between survey responses and provide additional internal validation of the survey.

**Table 4.**
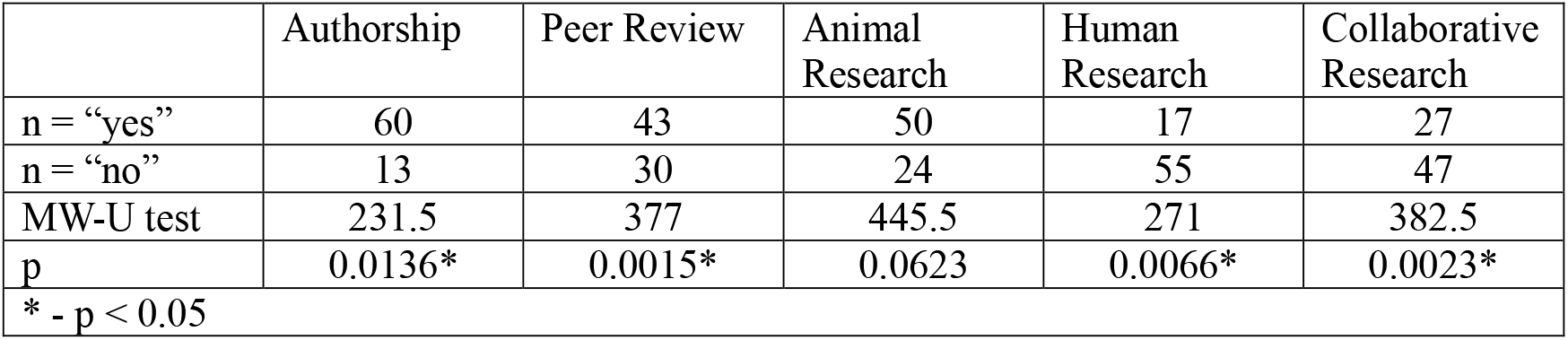
Differences in Responses to Survey Questions Between Groups Segregated Based Upon Experiences.

**Table 5.**
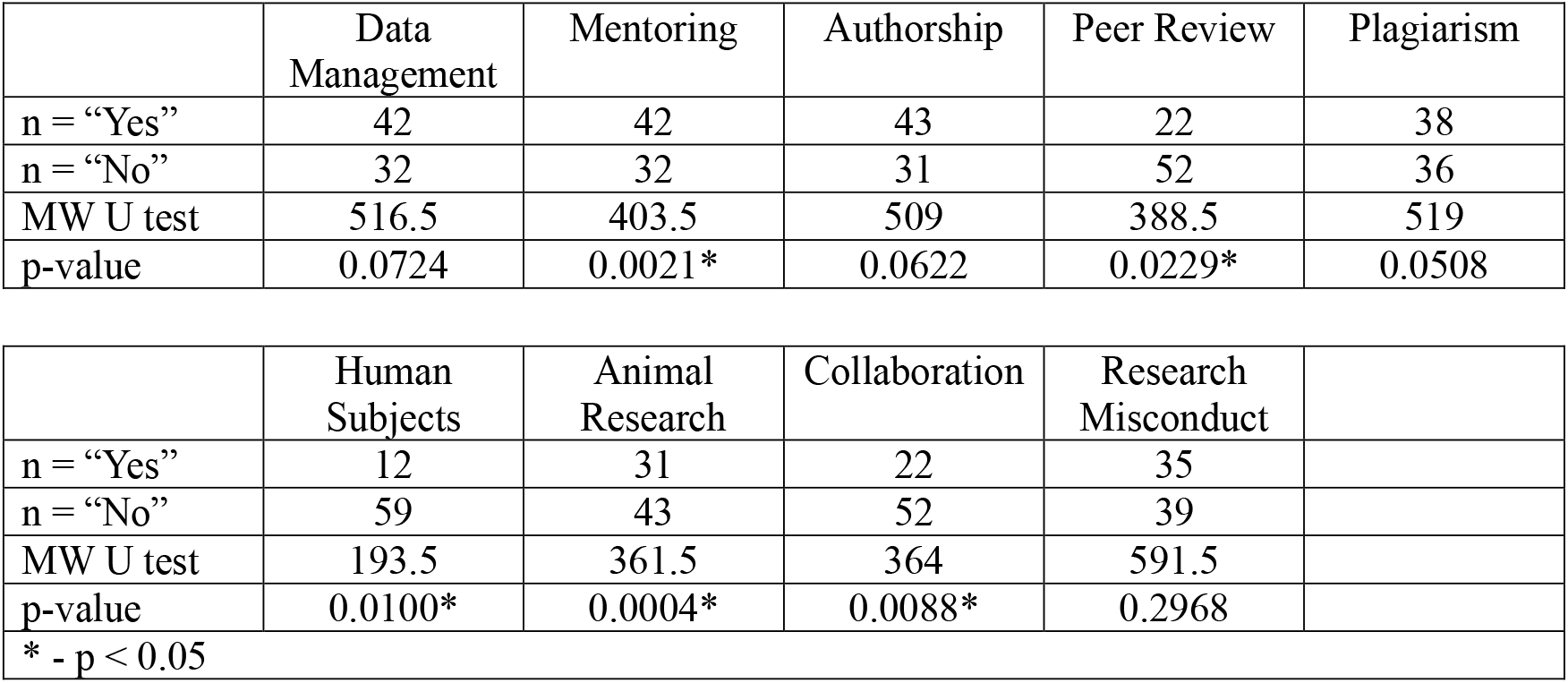
Differences in Response to Survey Questions Between Two Groups Segregated Based Upon Topics Indicated as Useful.

Finally, it was of interest to stratify the survey responses based upon progression towards completion of dissertation research to determine if overall responses changed with time in the graduate program. The responses to individual questions, e.g. usefulness of training in animal research, within different cohorts (Year 2, Year 3 or 4, Year 5 or 6, > Year 6 or Graduated), were compared with a Kruskal-Wallis test. The results show no statistically significant difference between these cohorts for any of the survey questions (**Online Resource 4**). The results suggest no differences in response based upon time in the graduate program. Stratifying the dataset resulted in some cohorts with small numbers and the outcome of this analysis may be limited by the sample size.

## Discussion

The results of the study reveal that the majority of students participate in training due to a mandate, however the motive for participation did not impact performance. Student attitudes were positive toward ethics training, and trainees recognize the importance of ethical issues in scientific research and the significance of training. The students learned from the course, and their self-efficacy increased. At the time of training, students could foresee the utility of training in different sessions. Retrospectively, past participants in the training found different elements of the course useful to the research in their doctoral studies. These findings suggest that from the student perspective, the significance of RCR training is recognized, and training is relevant to their research progression in graduate school. RCR training has had an impact on these trainees.

It was anticipated that students taking RCR training because it was mandated might have different attitudes, self-efficacy or knowledge gain than students taking the course for other reasons. Problems identified in RCR training include the requirement for RCR training not being taken seriously (M. Kalichman 2013a). Further, a meta-analysis of factors impacting RCR training outcomes identified mandating courses as a factor negatively impacting outcomes (Katsarov et al. 2022). Analysis of the outcomes of the current study reveals no differences in the outcomes measured – knowledge, attitudes or self-efficacy – between students taking the course due to a mandate and students providing a different rationale for taking the course. At least in this cohort of students, the motivation for taking the course had no impact on outcomes.

It was also anticipated that student attitudes towards research integrity and training might impact outcomes and that training might change the attitudes of students about RCR. At the beginning of the course, students strongly felt that consideration of ethical issues is important for scientific inquiry, that training is important and that training prevents misconduct. Due to the uniform positive attitude at the beginning of the course, it was not possible to stratify students based upon attitude to determine if attitude affected outcomes. Further, given the strong positive attitude at the beginning of the course, no differences in attitudes could be detected at the end of the course. Attitudes toward scientific integrity are important for reduced likelihood of scientific misconduct (Holm and Hofmann 2018). The goals of some RCR training includes improving the attitudes of trainees, and this change is considered an important outcome (Katsarov et al. 2022; McGee 2014). Given the important relationship of attitudes towards scientific integrity, outcomes of training and decreased occurrences of scientific misconduct, it was very encouraging to see the very positive attitudes of this cohort of students. These results are similar to other findings about student perceptions of RCR training (Langlais and Bent 2018).

Students indicated increased self-efficacy immediately following training in Fall 2023 and 2024. Believing in your own abilities and having confidence in your actions is essential to effectively navigate and solve ethical situations. However, inflated self-efficacy can be detrimental to RCR training and management of ethical situations. One hurdle to effectively engaging scientists in RCR training is their perception that they are completely capable of managing ethical situations already, which can be particularly true of graduate students (McCormick et al. 2012). Further, several studies have associated RCR training with increased unethical behaviors in some areas. This association might be related to inflated self-efficacy, and one study has correlated high self-efficacy in managing RCR with unethical behavior (Anderson et al. 2007; Antes et al. 2010). The increase in self-efficacy in the current study is encouraging especially since the retrospective reflections suggest that lessons learned from this training were useful in managing a range of issues related to RCR.

As described above, the evaluation of RCR training has focused upon knowledge and skills gained, and attitude and behavioral changes following training. Using these outcomes, recent studies have demonstrated a positive effect of training in scientific integrity (Watts et al. 2017; Abdi et al. 2021). Training appears to produce the outcomes believed to underlie scientific integrity by providing the foundation for trainees’ careers. It is also important that training is practical and relevant, and that trainees put lessons learned from training into practice to hone their skills (Nebeker 2014b, 2014a). The results from the current study demonstrate that trainees recognize the practicality of most RCR topics at the time of training. Similar to this finding, the results of a survey immediately following training indicated that 75% of trainees believe that the training program prepared them to avoid potential misconduct, recognize potential misconduct and respond to potential misconduct (Plemmons, Brody, and Kalichman 2006). The current study goes further to query past participants in the course (up to 7 years previous) about the utility of the course during their doctoral studies. The results of the survey of past participants in the RCR course reveal that many have utilized information provided during their training to navigate and/or avoid situations related to issues of scientific integrity. These results underscore the utility of RCR training for graduate students early in their career as it clearly prepares them to manage situations many will face as they proceed in their doctoral training and beyond.

Training in scientific integrity was implemented in response to numerous reports that led to policy changes at federal funding agencies, and training programs appear to successfully provide a foundation for performance of research with integrity. However, the implementation of training was only one recommendation in these reports, which emphasize creating a culture of scientific integrity at academic and research institutions (National Research Council 2002). It was recognized that training was a cornerstone of scientific integrity but was insufficient on its own if the culture of institutions promoted or allowed poor behavior or questionable scientific practices rather than scientific integrity. Institutional guidelines are required to clearly distinguish acceptable from unacceptable behavior. Appropriate actions must be fairly triggered in response to unacceptable behavior. High profile cases of academic misconduct provide the most egregious examples of misbehavior and while there are consequences, some argue that the penalties dispensed are insufficient (Devereaux 2014). In addition to providing regulations and policy to manage misconduct, institutions should also be responsible for modifying policies that provide the environment permissive for questionable scientific practices and misconduct in the first place. The “perverse incentives” that reward scientists and advance their careers provide the pressure that leads to the erosion of scientific integrity (National Research Council 2002; Steneck 2006; L. Bouter 2020). These incentives include the heavy reliance on quantitative metrics (numbers of publications, impact factor, citations) for evaluation for promotion and tenure, and the hypercompetitive environment for research funding and for research positions (L. Bouter 2020; Edwards and Roy 2017; L.M. Bouter 2015). Modification of policies to mitigate these incentives are necessary to truly create a culture of scientific integrity (Mejlgaard et al. 2020).

### Limitations

There are a number of limitations to this study. The sample size is relatively small, and the study was limited to two courses at a single site. The rate of consent for participation in the in-class study was high (94%), but the in-class surveys were reliant upon students faithfully reporting (remembering) a self-identified PIN for each survey (approximately 2 weeks apart). Eight sets of surveys were discarded since the post-surveys did not have PINs matching pre-survey PINs. Five additional students joined the class late and were not consented to participate. The response rate to the online survey was modest. While participation was incentivized, the incentive was small and offering a larger incentive might have yielded a better response rate. It is possible that responses to the online survey are not representative of the entire cohort and that students with particular views, e.g. positive views, of the impact of RCR training self-selected for participation in the study. It is also possible that the demographics of the class participants/survey respondents produced results that are not representative of the general graduate student population. For example, in an evaluation of ethical decision making of graduate students on issues related to scientific integrity, women and men performed significantly differently (Langlais and Bent 2014). Finally in retrospective analyses it is difficult to connect changes to specific interventions. The online survey was administered several years after training and many of the students had additional RCR training in the interim. The survey directed respondents to reflect on the impact of the BMS700 and/or BMS701 courses upon their ability to navigate issues of scientific integrity, but the influence of additional training cannot be excluded.

### Future Directions

This study serves as a pilot study of a single RCR training series at a single site. Future efforts should expand the study to increase the number of trainees, evaluate different training programs and survey trainees at different institutions. Expanded studies are necessary to determine the generalizability of the findings in this study for different cohorts of trainees at different institutions. These efforts will also provide insight into differences in student perceptions of RCR training at different types of institutions, e.g. at large private institutions vs small land grant universities. This study has addressed the perceptions of primarily graduate students about the utility of their RCR training experience, although a few postdoctorates were also included in the cohort. Since federal recommendations for scientific integrity include RCR training longitudinally in research careers, it would be informative to determine perceptions of the utility of RCR training at different career stages. One of the factors affecting the impact of RCR training and adult learning in general is the connection of the curriculum to the profession (Nebeker 2014b, 2014a). Studies addressing the perception of the utility of RCR training can directly assess connectedness to the scientific research profession and inform curricula modification to increase impact. Surveys at different career stages can provide insight into the development of longitudinal RCR training where distinct RCR topics are components of curricula designed for different career stages or where similar topics are explored to different depths depending on career stage. Efforts to increase utility of training along with incorporation of evidence-based types and methods of training (Mulhearn et al. 2017; Todd et al. 2017; Torrence et al. 2017) should direct development or modification of training programs in RCR.

## Supporting information

Supplemental Data - Online Resources

## Acknowledgements

The surveys were designed in consultation with France-Elvie Banda at FASEB (Federation of American Societies for Experimental Biology), Court Lanham at WVU, and Drs. Kelly Collins, Marieta Gencheva and Miriam Leary. Our thanks to a small group of students who participated in a focus group to validate the survey and improve its content.

## Notes

### Competing Interest Statement

The authors have declared no competing interest.

