## Supplemental Data - Online Resources for "Evaluation of the Utility of a Research Ethics Training Course to Graduate Students"

**Online Resource 1. Syllabi and Topics Covered in BMS700/BMS701**

**Scientific Integrity**
**BMS700 Fall Semester**

**Prerequisites:** None

**Course Time:** Every other Wednesday 3:00 to 4:50 PM

**Course Description:** As a trainee at the institution, you are required to meet particular federal and University-wide standards regarding the responsible conduct of research. This cours is one component of that along with online training via the Collaborative Institutional Training Initiative (CITI).
Note: BMS700 is the first of two semesters for training in the responsible conduct of research. In the Spring, BMS701, Scientific Rigor and Ethics will cover different topics related to the responsible conduct of research.

**Format:** Part didactic, part discussion-based

**Required Textbooks:** Steneck, 2006, ORI Introduction to the Responsible Conduct of Research Available for download: https://ori.hhs.gov/ori-intro

**Course Objectives:** In completing the fall semester course, the trainees will be able to:
1. Describe the primary issues that represent breaches in scientific integrity
2. Describe the context that can result in poor ethical decision making
3. Describe the steps to take if you believe a breach of scientific integrity has occurred
4. Define plagiarism and identify different forms of plagiarism. Describe strategies to prevent plagiarism in your own writing
5. Collect data in a manner consistent with the standards of funding agencies and with proper research approaches
6. Identify the activities that represent responsible data collection with regard to images. Describe the steps where image manipulation is acceptable
7. Define what are appropriate social interactions and describe what you need to do to consider the rights of others with whom you interact during your career
8. Describe the steps needed to use animals in research and identify the responsibilities of laboratory personnel in the use of animals
9. Describe the steps needed to use humans in research and identify the responsibilities and best practices for laboratory personnel in the use of human subjects

**Grading:** The grade reflects the results of two exams, attendance and class participation. Exams are take-home format. You may use the text and your notes, but you may not discuss with other students in the course. Absences from the course require prior approval by the instructors. More than one unexcused absence will result in a grade of "Incomplete" and the trainee will need to make up the course the subsequent year

**Grading policy:** The course is graded P/F based on exams, attendance and participation in case discussions. A grade of P requires a combined score of 75% or better on the two exams and attendance of all sessions for the entire time of the session. Participation in discussions will reflect active exchanging of information within the small group.

**Course Schedule:**
Week 1 – Plagiarism
Week 2 – Social Responsibility
Week 3 – Recording Research Data
Week 4 – Reporting Photographic Research Results
Week 5 – Ethical Decision Making – The Lab exercise
Week 6 – Human Research Protections
Week 7 – Animal Care and Use

### Scientific Rigor and Ethics
### BMS701 Spring Semester

**Prerequisites:** None. It is not necessary to complete BMS700 prior to enrollment in BMS701

**Course Time:** Every other Wednesday 3:00 to 4:50 PM

**Course Description:** As a trainee at the institution, you are required to meet particular federal and University-wide standards regarding the responsible conduct of research. This course is one component of that along with online training via the Collaborative Institutional Training Initiative (CITI).
Note: BMS701 is the second of two semesters for training in the responsible conduct of research. In the Fall, BMS700, Scientific Integrity covers different topics related to the responsible conduct of research.

**Format:** Part didactic, part discussion-based

**Required Textbooks:** Steneck, 2006, ORI Introduction to the Responsible Conduct of Research
Available for download: https://ori.hhs.gov/ori-intro

**Course Objectives:** In completing the spring semester course, the trainees will be able to:
1. Become aware of challenges that women in science must overcome to succeed
2. Define conflicts of interest. Describe how to identify conflicts of interest
3. Discuss the role of the scientist in today's society, including explaining science to non-scientists and advocating for good scientific research.

4. Define criteria that can result in authorship. Describe the peer review process.
5. Define collaboration. Describe the benefits of collaboration to research. Identify signs that indicate good practice has been breached in collaboration. List steps to develop a strong and effective collaboration.
6. List the issues for when considering sex in designing experiments is necessary for a rigorous test of the experimental question.
7. Describe problems with rigor and reproducibility and solutions to increase rigor.

**Grading:** The grade reflects the results of two exams, attendance and class participation. Exams are take-home format. You may use the text and your notes, but you may not discuss with other students in the course. Absences from the course require prior approval by the instructors. More than one unexcused absence will result in a grade of "Incomplete" and the trainee will need to make up the course the subsequent year

**Grading policy:** The course is graded P/F based on exams, attendance and participation in case discussions. A grade of P requires a combined score of 75% or better on the two exams and attendance of all sessions for the entire time of the session. Participation in discussions will reflect active exchanging of information within the small group.

**Course Schedule:**
Week 1 – Conflict of Interest
Week 2 – Authorship and Peer Review
Week 3 – Collaboration
Week 4 – "Picture a Scientist"
Week 5 – The Scientist in Society
Week 6 – Sex as a Biological Variable
Week 7 – Rigor and Reproducibility

**Online Resource 2. Surveys used in the Study**

**Pre-Survey used in Fall 2023 and Fall 2024**

**Attitudes**
Question 1
The primary reason I am taking this course is because _____________.
a. it is mandated.
b. I am interested in ethical issues related to science.
c. it is my responsibility as a scientist to be trained in ethics.

Question 2
Ethical issues are important to consider in the conduct of scientific inquiry.

a. Strongly disagree
b. Disagree
c. Neither agree nor disagree
d. Agree
e. Strongly agree

Question 3
Training in scientific integrity/ethics is very important for scientists.
a. Strongly disagree
b. Disagree
c. Neither agree nor disagree
d. Agree
e. Strongly agree

Question 4
Training in scientific ethics can help prevent research misconduct.
a. Strongly disagree
b. Disagree
c. Neither agree nor disagree
d. Agree
e. Strongly agree

Question 5
How frequently do ethical issues in science arise?
a. Never
b. Hardly ever
c. Occasionally
d. Frequently

**Self-Efficacy**
Question 6
How knowledgeable are you about ethical issues in science?
a. I have no knowledge
b. I have a little knowledge
c. I have about average knowledge
d. I have more than average knowledge
e. I am very knowledgeable

Question 7
How confident are you in your ability to handle ethical issues in science?
a. I have no confidence
b. I have some confidence
c. I have a lot of confidence
d. I am very confident

Question 8
How confident are you in your ability to find the resources necessary to handle ethical issues in science?
a. I have no confidence
b. I have some confidence
c. I have a lot of confidence
d. I am very confident

Question 9
How confident are you in your ability to maintain ethical behavior in challenging situations?
a. I have no confidence
b. I have some confidence
c. I have a lot of confidence
d. I am very confident

Question 10
I can apply ethical principles and standards to guide decision making.
a. Strongly disagree
b. Disagree
c. Neither agree nor disagree
d. Agree
e. Strongly agree

**Knowledge**

1. Using your own written material for more than one purpose is-
   a) An example of self-plagiarism and acceptable
   b) Acceptable when referenced in a footnote where else the material is used
   c) Acceptable only when the material is unpublished
   d) Unacceptable due to self-plagiarism
   e) Unacceptable due to federal law

2. What is the best way to prevent unconsciously plagiarizing from a document?
   a) Make a photocopy of original document
   b) Read the original document upside down so it is harder to recall
   c) Take notes while reading the document and refer only to those notes
   d) Copy-and-paste from the document with highlighting to remember what was copied
   e) Use quotation marks around anything that might be close to plagiarized

3. The best way to avoid offending others in your work environment is to-
   a) Only say things that you don’t find offensive
   b) Avoid saying anything personal to stay out of trouble
   c) Think how others might interpret what you have said
   d) Say whatever comes to your mind and then ask others if it was offensive

4. A microaggression is-
a) A small cluster of aggregated proteins
b) A conflict that results in pushing and shoving but no blows being thrown
c) Usually considered as sexual harassment
d) A form of subtle or unintentional discrimination

5. A laboratory notebook is an important record of your experiments. What piece of information is NOT likely to be helpful for later repetition of your experiments?
a) The date the experiment is done
b) The time of day that you run your gel
c) The temperature in the lab (for room-temperature incubations)
d) The sex of the cell line used for cell culture
e) The primer sequences used for a PCR reaction

6. When you finish your dissertation, what happens to your lab notebooks?
a) They should be shredded to protect your intellectual property rights
b) They are retained by you for reference and safe keeping
c) They are retained by the WVU library as reference material
d) They are retained by your mentor for safe keeping
e) They are retained by the WVU Research Office for compliance purposes

7. When working with confocal image data, which of these changes are not appropriate?
a) Changing the image size to save disk space
b) Changing the gamma setting
c) Changing the contrast setting
d) Changing the brightness setting
e) Rotating the image 90 degrees

8. What is the optimal format to save a scientific image?
a) JPG
b) Powerpoint
c) TIFF
d) Word
e) GIF

9. If you experience an ethical dilemma during your graduate education, who would be the least biased person to contact to report the incident?
a) Your principal investigator
b) A fellow lab mate
c) The university's provost
d) The university's research integrity officer
e) The department's chair

10. Reporting ethical misconduct to authorities-
a) Absolves you of any responsibility
b) Has no consequences on your research project
c) Prevents your identity from being determined
d) Protects others in your lab from scrutiny
e) Could lead to retaliation against you

11. Which of the following principles is not normally considered when work with human research subjects?
a) Regional representation
b) Beneficence
c) Respect for persons
d) Justice

12. Many of the principles of working with human subjects arose from atrocities that occurred throughout the years. These rules have been established by all BUT which of the following groups?
a) The Nuremberg Code of 1947
b) The Geneva Convention of 1949
c) The Helsinki Declaration of 1964
d) American Psychological Association Guidelines of 1973
e) The Guidelines by the U.S. Department of Health, Education, and Welfare Codes of 1974

13. Three principles (the 3 R's) are important considerations for working with animals in research. Which of the following is not one of the 3 R's?
a) Reduction
b) Refinement
c) Reuse
d) Replacement

14. Which agency has jurisdiction over the use of animals that are used in biomedical research in the US (except mouse, rats, and birds that are bred for research)?
a) US Department of Agriculture
b) US Food and Drug Administration
c) US Department of Commerce
d) UN World Food Program
e) US Justice Department

**Post-Survey used in Fall 2023 and Fall 2024**

The questions on the post-survey were identical to the questions on the pre-survey except for one question. The question from the pre-survey:

The primary reason I am taking this course is because _____________.
a. it is mandated.
b. I am interested in ethical issues related to science.
c. it is my responsibility as a scientist to be trained in ethics.

was replaced with the question:

This course has increased my confidence in my ability to handle ethical issues in science
a. Strongly disagree
b. Disagree
c. Neither agree nor disagree
d. Agree
e. Strongly agree

**Demographics Survey used in Fall 2024 and Online**

With which gender do you most identify? *Select all that apply to you.*

- o Non-binary
- o Transgender woman
- o Cisgender woman
- o Transgender man
- o Cisgender man
- o Prefer to self-describe
- o Prefer not to answer

If you prefer to self-describe above, please do so here ____________________

An individual's racial and ethnic identity is their sense of belonging or connectedness to a group who shares their culture and socialized racial-ethnic experience. How do you identify yourself in terms of race and ethnicity? *Select all groups that apply to you.*

- o Hispanic/Latino/a/x – a person of Cuban, Mexican, Puerto Rican, South or Central American, or other Spanish culture or origin, regardless of race.
- o Not Hispanic/Latino/a/x
- o Indigenous Person, American Indian or Alaska Native – a person having origins in any of the original peoples of North or South America (including Central America), and who maintains tribal affiliation or community attachment
- o Asian – a person having origins in any of the original peoples of the Far East, Southeast Asia, or the Indian subcontinent, including, for example, Cambodia, China, India, Japan, Korea, Malaysia, Pakistan, the Philippine Islands, Thailand, or Vietnam.

- o Black or African American – a person having origins in any of the Black racial groups of Africa.
- o Native Hawaiian or Other Pacific Islander – a person having origins in any of the original peoples of Hawaii, Guam, Samoa or other Pacific islands.
- o Middle Eastern or North African – a person having origins in any of the original peoples of Southwest Asia and North Africa for example: Iran, Lebanon, Syria, Yemen, Libya, Morocco, or Algeria
- o White – a person having origins in any of the original peoples of Europe.
- o Prefer to self-describe
- o Prefer not to answer

If you prefer to self-describe above, please do so here ___________________

Are you the first-generation in your immediate family to pursue/obtain a post-secondary degree in the U.S. or abroad?

- o Yes
- o No
- o Prefer not to answer

Do you self-identify as a person with a disability (physical or mental health)?

- o Yes
- o No
- o Prefer not to answer

Did you grow up in or do you currently live in a low-income household? *Please select all that apply*

Please note: Current thresholds of low-income households are $20,385 for a single-member household and $69,945 for an eight-member household.

- o Yes, I grew up in a low-income household
- o No, I did not grow up in a low-income household
- o Yes, I currently live in a low-income household
- o No, I do not currently live in a low-income household
- o Prefer not to answer

I am a domestic/international student

- o I am a domestic (US) student

- I am an international student
- Prefer not to answer

**Online Scientific Integrity Training Survey Used for Past Participants in the Course**

What year in your graduate studies have you completed?

- Year 2 or 3
- Year 4 or 5
- Year 6 or above
- Graduated already
- Prefer not to answer

What type of work have you done for your dissertation studies? *Check all that apply*

- Human subject research
- Animal research
- In vitro studies
- Computational studies
- Collaborative research
- Health policy
- Other (please specify)
- Prefer not to answer

If you answered Other above, please specify here ________________

Have you published during your time at WVU?

- Yes
- No
- Submitted manuscript but not yet published
- Prefer not to answer

Have you participated in peer review of a manuscript while at WVU?

- Yes
- No
- Prefer not to answer

Besides the BMS700/BMS701 Scientific Integrity courses, have you had any additional training in the Responsible Conduct of Research (RCR)?

- Yes
- No
- Prefer not to answer

What additional training in RCR have you had? ____________________

Guidance for answering questions:

Consider the example question “I have used the concepts from the course in discussions to manage the development of intellectual property.”

Possible answers are: strongly agree, agree, neither agree nor disagree, disagree, strongly disagree

What if you have not been involved in the development of intellectual property thus far in your career? You should choose the answer “strongly disagree”. Even though you might have learned important concepts from the course that you intend to apply in future, you have not yet used the concepts in this situation.

Consider the content and experiences from the BMS700/BMS701 Scientific Integrity courses and how these have helped you navigate specific situations.

I have used lessons from one or more sessions of the course for managing my lab notebook or data collection.

- Strongly disagree
- Disagree
- Neutral
- Agree
- Strongly agree

I have used concepts from the course to develop my relationship with my mentor.

- Strongly disagree
- Disagree
- Neutral
- Agree
- Strongly agree

I have used concepts from the course to fulfill my responsibilities as an author/co-author.

- Strongly disagree

- o Disagree
- o Neutral
- o Agree
- o Strongly agree

I have used knowledge from the session on peer review to guide my critique of manuscripts.

- o Strongly disagree
- o Disagree
- o Neutral
- o Agree
- o Strongly agree

The plagiarism session did **NOT** provide useful examples to avoid ethical issues.

- o Strongly disagree
- o Disagree
- o Neutral
- o Agree
- o Strongly agree

I have applied concepts from the course to make sure that my experiments on human subjects are performed ethically?

- o Strongly disagree
- o Disagree
- o Neutral
- o Agree
- o Strongly agree

I have applied concepts from the course to make sure that my experiments on animals are performed ethically.

- o Strongly disagree
- o Disagree
- o Neutral
- o Agree
- o Strongly agree

I have applied concepts from the collaborative science session to improve interactions with collaborators.

- o Strongly disagree

- Disagree
- Neutral
- Agree
- Strongly agree

Training from the course provided knowledge to manage research misconduct situations.

- Strongly disagree
- Disagree
- Neutral
- Agree
- Strongly agree

Have you managed a conflict of interest situation during your studies?

- Yes
- No
- Prefer not to answer

If student's answered Yes to the previous question, they were prompted with the following question-

I used knowledge from the course to manage a conflict of interest situation.

- Strongly disagree
- Disagree
- Neutral
- Agree
- Strongly agree

During your time in graduate school, have you experienced an ethical dilemma where the content or experience from the ethics courses was useful to help you navigate the situation? What specific topics or scenarios were covered that you found relevant to the situation? *Check all that apply*

- Data management/ownership
- Conflict of interest
- Human subjects
- Animal welfare
- Research misconduct
- Plagiarism
- Publication and authorship
- Mentor/trainee responsibilities
- Peer review

- Collaborative science
- I have not experienced an ethical dilemma where the ethics course material helped

Were there any topics that were taught in the ethics courses that you found useful to help avoid unethical situations? What specific topics or scenarios were covered that you found relevant to the situation? *Check all that apply*

- Data management/ownership
- Conflict of interest
- Human subjects
- Animal welfare
- Research misconduct
- Plagiarism
- Publication and authorship
- Mentor/trainee responsibilities
- Peer review
- Collaborative science
- I have not experienced unethical situations that the ethics course material helped avoid

Were there ethical issues that you have experienced where the course content was not sufficient to manage the situation or was not covered in the course? If so, please specify the topic.

**Online Resource 3. Motivation to Take RCR Training Does Not Impact Outcomes**

| | **Test** | **t or Mann Whitney U** | **p value** |
|---|---|---|---|
| **Knowledge – pre** | t-test | 1.767 | 0.0810 |
| **Knowledge – post** | t-test | 1.622 | 0.1086 |
| **Knowledge - delta** | t-test | 0.1397 | 0.8892 |
| **Self-Efficacy – pre** | t-test | 1.705 | 0.0920 |
| **Self-Efficacy – post** | t-test | 0.5960 | 0.5960 |
| **Self-Efficacy - delta** | t-test | 1.009 | 0.3161 |
| **Training is important** | Mann Whitney | 706.5 | 0.0818 |
| **Training prevents misconduct** | Mann Whitney | 831.5 | 0.87 |
| **Ethical issues are important in science** | Mann Whitney | 228 | 0.3383 |

Comparison of two cohorts: students taking the course due to mandate vs. students taking the course for other reasons (because it's their responsibility or out of interest). The responses of the two cohorts to knowledge questions (pre or post-test) and self-efficacy questions (pre or post-test) were analyzed. The sum of the scores of the self-efficacy questions was used in the analysis. The difference between their scores on the post-test and the pre-test (delta) was also analyzed. There was no significant difference in response to attitude questions on the pre- and post-test. The responses to the questions on the pre-test were compared between cohorts. As the data is incomplete due to a survey error, the responses to each attitude question rather than the sum of responses was compared between the two cohorts. Knowledge and self-efficacy summed scores were Gaussian and were analyzed using a t-test. Attitude responses were not Gaussian so a nonparametric Mann Whitney test was performed. All tests performed were unpaired, two-tailed.

**Online Resource 4. Results of Online Survey Stratified by Year in Grad School.**

| | n | KW Statistic | p-value |
|---|---|---|---|
| Data Management | 73 | 1.956 | 0.7439 |
| Mentoring | 73 | 1.214 | 0.8758 |
| Authorship | 73 | 1.637 | 0.8022 |
| Peer Review | 73 | 2.456 | 0.6526 |
| Plagiarism | 73 | 8.104 | 0.0878 |
| Human Subjects | 71 | 2.145 | 0.7092 |
| Animal Research | 73 | 1.445 | 0.8364 |
| Collaboration | 73 | 1.329 | 0.8565 |
| Research Misconduct | 73 | 7.505 | 0.1115 |

The responses to the questions related to these nine sessions were stratified based upon progression of the respondent through graduate school, e.g. in Year 3 or 4. The responses of each of the resulting cohorts were compared using a Kruskal-Wallis test.
